# Positive selection and enhancer evolution shaped lifespan and body mass in great apes

**DOI:** 10.1101/2021.07.08.451631

**Authors:** Daniela Tejada-Martinez, Roberto A. Avelar, Inês Lopes, Bruce Zhang, Guy Novoa, João Pedro de Magalhães, Marco Trizzino

## Abstract

Within primates, the great apes are outliers both in terms of body size and lifespan, since they include the largest and longest-lived species in the order. Yet, the molecular bases underlying such features are poorly understood. Here, we leveraged an integrated approach to investigate multiple sources of molecular variation across primates, focusing on ~1,550 genes previously described as tumor suppressors, oncogenes, ageing genes in addition to a novel Build of the CellAge database of cell-senescence genes (version 2), herein presented for the first time. Specifically, we analyzed dN/dS rates, positive selection, gene expression (RNA-seq) and gene regulation (ChIP-seq). By analyzing the correlation between dN/dS, maximum lifespan and body mass we identified 67 genes that in primates co-evolved with those traits. Further, we identified 6 genes, important for immunity, neurodevelopment and telomere maintenance (including *TERF2*), under positive selection in the great ape ancestor. RNA-seq data, generated from the liver of six species representing all the primate lineages, revealed that ~8% of the longevity genes are differentially expressed in apes relative to other primates. Importantly, by integrating RNA-seq with ChIP-seq for H3K27ac (which marks active enhancers), we show that the differentially expressed longevity genes are significantly more likely than expected to be located near a novel “ape-specific” enhancer. Moreover, these particular ape-specific enhancers are enriched for young transposable elements, and specifically SINE-*Vntr*-Alus (SVAs). In summary, we demonstrate that multiple evolutionary forces have contributed to the evolution of lifespan and body size in primates.

## Introduction

Uncovering the molecular and evolutionary bases of ageing in the tree of life is important to increase our understanding of the natural mechanisms of disease resistance (Galis and Metz 2003; Caulin and Maley 2011; Ganten and Nesse 2012; Nesse et al. 2012; Thomas et al. 2013; Petralia et al. 2014; Abegglen et al. 2015; Harris et al. 2017; Ciccarelli and DeGregori 2020). The evolution of lifespan in animals has been shaped by selective pressures arising from different environments, ecological niches, habitats, and diets (Healy et al. 2014; Tollis et al. 2017; Kacprzyk et al. 2021). Ageing and senescence are the result of gradual declines in biological functions, which lead to increased vulnerability, disease predisposition, and ultimately death (López-Otín et al. 2013). With the exception of a few species that show no signs of ageing (e.g. some jellyfishes and hydras; (Petralia et al. 2014), most animals undergo this process. Nevertheless, maximum lifespan is a highly variable trait across the tree of life, suggesting that long-lived species may have adopted several evolutionary strategies which may have favored longevity by reducing disease susceptibility (Gorbunova et al. 2014; Kacprzyk et al. 2021; Yu et al. 2021). Yet, most of these strategies remain unknown.

One of the main consequences of ageing is the increased risk of developing cancer. Cancer has been reported in almost all multicellular organisms (Aktipis et al. 2013; Albuquerque et al. 2018), and it is often associated with somatic genetic mutations that inactivate tumor suppressor genes or activate oncogenes. These mutations ultimately affect cell metabolism, promoting uncontrolled cell division (Stratton et al. 2009; Tomasetti and Vogelstein 2015). Statistically, animal species with larger body sizes and longer lifespans accumulate more cell divisions during their lives and are therefore expected to accumulate more deleterious genetic mutations. Nonetheless, large and long-lived species overall do not develop cancer at higher frequency than smaller species (Nunney 2018; Seluanov et al. 2018). This scientific paradox was first noted by Richard Peto (the “Peto’s Paradox”; Peto et al. 1977; Nunney 1999).

Specific anti-cancer mechanisms have been discovered in long-lived species. These mechanisms include early contact inhibition in naked mole rats (Seluanov et al. 2009; Tian et al. 2013), positive selection in DNA repair & inflammatory genes in the giant tortoises from Galapagos (Quesada et al. 2019), and immunity pathways, telomere maintenance and cellular senescence (CS) genes in bats (Zhang et al. 2013; Foley et al. 2018; Tian et al. 2018; Wilkinson and Adams 2019). Particularly, CS - a process in which otherwise replicating cells reach the maximum number of divisions and cease proliferating - is another important mechanism of cancer suppression; previous studies indicate that genes regulating the senescent phenotype are strongly conserved amongst vertebrates compared to other protein-coding genes (Avelar et al. 2020). Despite the importance of senescence processes in inhibiting cancer, CS itself is detrimental for health; senescent cells produce pro-inflammatory cytokines that can paradoxically promote cancer (Rao and Jackson 2016). A previous study found that senolytics - drugs that specifically target and eliminate senescent cells - can enhance life- and health-span in old mice (Xu et al. 2018). On another hand, in larger mammals such as elephants and whales, molecular substitutions in cell senescence genes and increases in the copy number of important tumor suppressor genes have contributed to significantly reduce the risk of developing cancer (Abegglen et al. 2015; Caulin et al. 2015; Keane et al. 2015; Tollis et al. 2019). Specific changes in the regulation and expression of specific genes have also been associated with ageing (de Magalhães et al. 2009; Donlon et al. 2017; Morris et al. 2019; Chatsirisupachai et al. 2021). Yet, the extent of the contribution of cis-regulatory evolution to ageing is still poorly understood.

In primates, maximum lifespan (MLS) and body mass (BM) are correlated, despite being highly variable across species, with great apes (gorillas, orangutans, humans and chimpanzees) representing outliers (Finch and Austad 2012). The BM ranges from 5 kg in the grey mouse lemur (*Microcebus murinus*) to ~140kg in the gorilla (*Gorilla gorilla*). Similarly, the MLS varies from 13 years in the calabar angwantibo (*Arctocebus calabarensis*) to 55–122 years in great apes (AnAge database: https://genomics.senescence.info/; Tacutu et al. 2018). Outside great apes, just a few primate species can reach 50 years (e.g. the capuchin monkeys; Muntané et al. 2018; Orkin et al. 2021).

Consistent with Peto’s Paradox, great apes develop cancer at lower rates than other primates (Fowler et al. 1980; Lowenstine 1986; Cho et al. 2007; Lowenstine et al. 2016). Humans represent an exception, possibly as a consequence of their life-style (Finch 2010; Hochberg and Noble 2017; Albuquerque et al. 2018). In fact, in some human populations cancer rates are ~25% (Ferlay et al. 2015), and some cancer types seem to be unique to our species (e.g prostate cancer and lung cancer). Several studies have recently investigated the link between longevity, body mass and disease resistance in several mammalian species (Tollis et al. 2017; Muntané et al. 2018; Boddy et al. 2020; Huang et al. 2021; Kacprzyk et al. 2021). On the other hand, the mechanisms of ageing in great apes are largely uncharacterized, and several questions remain unanswered.

Here, we aimed at investigating the relative contribution of different sources of molecular variation (molecular evolution in coding genes, positive selection, cis-regulatory and gene expression evolution) to the evolution of longevity in great apes. First, we performed a literature search for genes that are involved in the CS process. This led to the generation of a novel Build (version 2) of the CellAge database, which includes 934 genes that induce or inhibit cellular senescence *in vitro* in human cell lines following genetic manipulation (Table S2).

Next, we interrogated 19 mammalian genomes, focusing on ~1,550 longevity-related genes, including the genes in the novel CellAge database, plus hundreds of genes previously associated with tumor suppression, oncogenic activity and ageing-related genes processes. We evaluated: 1) potential correlations between MLS, BM and coding gene evolution [non-synonymous/synonymous mutations]; 2) positive selection signal in the great-ape ancestor; 3) species-specific gene expression patterns (RNA-seq); 4) species-specific gene-regulation patterns (ChIP-seq for H3 lysine 27 acetylation [H3K27ac], which marks active cis-regulatory elements).

We identified 67 longevity genes that co-evolved with lifespan and/or body mass in primates, and 6 longevity genes under positive selection in the great-ape ancestor. These genes are important for immunity, neurodevelopment and telomere maintenance. In addition, RNA-seq and ChIP-seq data, generated from the liver of six species representing all the primate lineages (Trizzino et al. 2017), revealed that ~8% of the longevity genes investigated in this study are differentially expressed in apes relative to other primates, and that these lineage-specific gene expression patterns are associated with the rise of novel ape-specific enhancers, most of which derived from the insertion of young transposable elements.

## Results

### Ape-specific patterns in the evolution of lifespan and body mass

The great apes are the primates with the largest body size, and are also the most long-lived species in the entire order (fig. 1a-b). Given the high diversity of phenotypes across primates, we used Phylogenetic Generalized Least Squares (PGLS) models (Orme et al. 2012; Revell 2012; Pennell et al. 2014) to evaluate if body mass (BM) and maximum lifespan (MLS) evolved independently in great apes. The PGLS estimate the covariance among traits, MLS and BM in this case, taking into account the effect of the phylogenetic signal across the tree. We found that the allometric expectations are maintained across primates, with a positive correlation between BM and MLS (fig 1a). Our model revealed that the BM predicts ~6% of the variation in longevity across primate species (adjusted R-squared: 0.05904, 131 degrees of freedom, *p* = 0.0028). Nonetheless, the relation between MLS and BM in great apes is significantly different from the other primates (adjusted R-squared: 0.04568, degrees of freedom, *p* = 0.02888). This suggests that lineage specific molecular evolution may have shaped lifespan and body mass in great apes.

**Figure 1.**
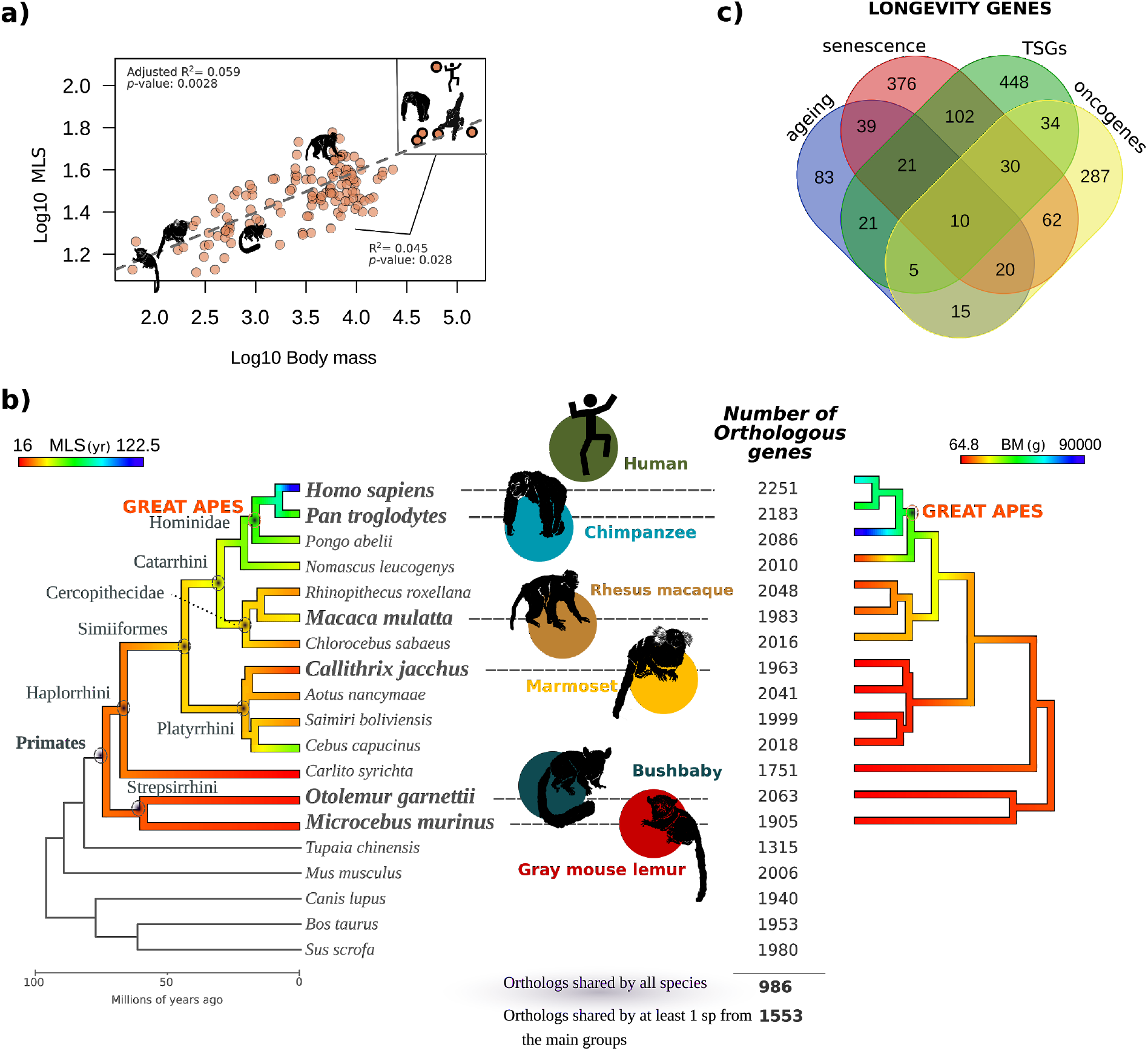
The evolution of body mass (BM) and maximum lifespan (MLS) across primates. a) Phylogenetic generalized least-squares models (PGLS) correlating MLS ~ BM across primates. The dashed lines represent a positive correlation between log10-transformed BM and MLS. The continuous line displays the correlation between great apes and other primates. b) Phylogenomic design. The molecular evolution analysis included 19 mammalian species. The six species highlighted are representatives from the main primates lineages (Catarrhini, Platyrrhini and Strepsirrhini). For these six species, RNA-seq and ChIP-seq data were publicly available (Trizzino et al. 2017). The colours in the phylogenetic tree reflect the values of MLS and BM in each primate branch respectively, from lowest (red) to highest (blue). c) Venn diagram representing the total number of orthologous longevity genes (1,553) shared by at least one species in each of main primates lineages, broken down based on the different categories (tumor suppressors, oncogenes, cell senescence and ageing genes).

### Build 2 of the CellAge Database of Cell Senescence Genes

We compiled a Build 2 of the CellAge dataset by means of a scientific literature search of gene manipulation experiments in primary, immortalized, or cancer human cell lines that caused cells to induce or inhibit CS. The novel CellAge build comprises 934 distinct CS genes, of which 608 genes affect replicative CS (our default annotation), 180 genes affect stress-induced CS, and 214 genes are related to oncogene-induced CS. Of the 934 total genes, 393 genes induce CS (~ 42.1%), 518 inhibit it (~ 55.5%), and 23 genes have unclear effects, both inducing and inhibiting CS depending on experimental conditions (~ 2.5%). The genes in the dataset are also classified according to the experimental context used to determine these associations (Table S2).

### The rate of evolution (dN/dS) in 67 genes positively correlates with the evolution of longevity and body mass across primates

The ratio between synonymous (dS) and nonsynonymous substitutions (dN) is considered a reliable measure of natural selection in the evolution of the phenotypic diversity. In line with this, several studies have reported a positive relationship between dN/dS and either body mass or maximum lifespan (Romiguier et al. 2010; Weber et al. 2014; Figuet et al. 2016). Nonetheless, the contribution of individual genes to the evolution of body size and lifespan remains largely unexplored, especially in primates. Even if MLS and BM are positively correlated in most mammalian lineages, their molecular evolution is likely largely independent, as has been previously reported in rodents (Seluanov et al. 2018). To investigate the molecular evolution of longevity and body mass in primates, we focused on a set of 2,268 genes previously implicated with lifespan, body mass and cancer (hereafter “longevity genes”). This list includes genes that have been previously associated with tumor suppression, with oncogenic functions, and with ageing and senescence-related processes (Table S1.2). Of these 2,268 genes, 1,553 have a six-way ortholog in six species (human, chimpanzee, rhesus macaque, marmoset, bushbaby, mouse lemur), representative of the main primate lineages (Catarrhini, Platyrrhini and Strepsirrhine), for which we have also analyzed publicly available next generation sequencing data (see below). These 1,553 genes were retained for downstream analyses. Additionally, 986/1,553 (~63.5%) of these orthologous genes are also present in 11 additional primate species that we included in our molecular evolution analysis (fig. 1b-c, Table S1.3). In general, the number of orthologous genes was similar across primates, with a variation between 1,905 genes in the bushbaby to 2,183 in the chimpanzee. The species with the smallest number of orthologous genes was the tarsier (*Carlito syrichta*), likely reflecting the low sequencing coverage for the genome of this species (Table S1.1).

We correlated the rates of evolution (dN/dS) in the 986 genes with the two life history traits (body mass [BM], and maximum lifespan [MLS]) together and independently (Table S1.4-1.6). As a first step, we removed 89 genes with ω>2 due to the possible overestimation of ω in the branch due to saturation or miss calculation of synonymous (dN) or non-synonymous (dN) substitutions. Next, we used the variance of the residuals from the first PGLS model (MLS ~ BM), in order to account for the expected covariance structure between the two variables and evaluate the contribution of each gene (dN/dS) in the evolution of MLS and BM simultaneously. As a result, we found 9 genes for which the dN/dS is positively correlated with both MLS and BM (figure 2a, table 1): five are tumor suppressor genes (TSGs: *PRDM2 , ADAMTS8 , NPRL2 , SGMS1 , LRIG1*), one is an TSG-oncogene (*NCOA4*), one is categorized as TSG-oncogene-senescence gene (*ATF3*) and one is an ageing gene (*TRAP1*). These genes are involved in important biological pathways. For instance, *PRDM2* is involved in cell differentiation (Fog et al. 2012) and *NCOA4* in iron signaling, which is a key autophagic cell death process (Bellelli et al. 2016). Notably, *ADAMTS8* contributes to ~30% of the variation in MLS and BM across primates. This gene encodes for a protein involved in several types of cancer (Zhao et al. 2018), it is important for the immune response (Zhang et al. 2020) and has been associated with multiple diseases such as rheumatoid arthritis (Baloğlu et al. 2020). Importantly, *ADAMTS8* has also been reported as a positively selected gene in whales, the longest-lived/most cancer-resistant group of species across mammals (Tejada-Martinez et al. 2021). Gene enrichment analysis on the nine genes (*p*<0.05, Table 1, Table S1.7; complete list of longevity genes used as background) revealed an overrepresentation in ontology process related with response to amino acid starvation (*NPRL2* and *ATF3*), which could reflect the important role that the diet plays in adaptation, longevity and species diversification (Yabu et al. 2012; Gallinetti et al. 2013; Slade and Staveley 2016).

**Figure 2.**
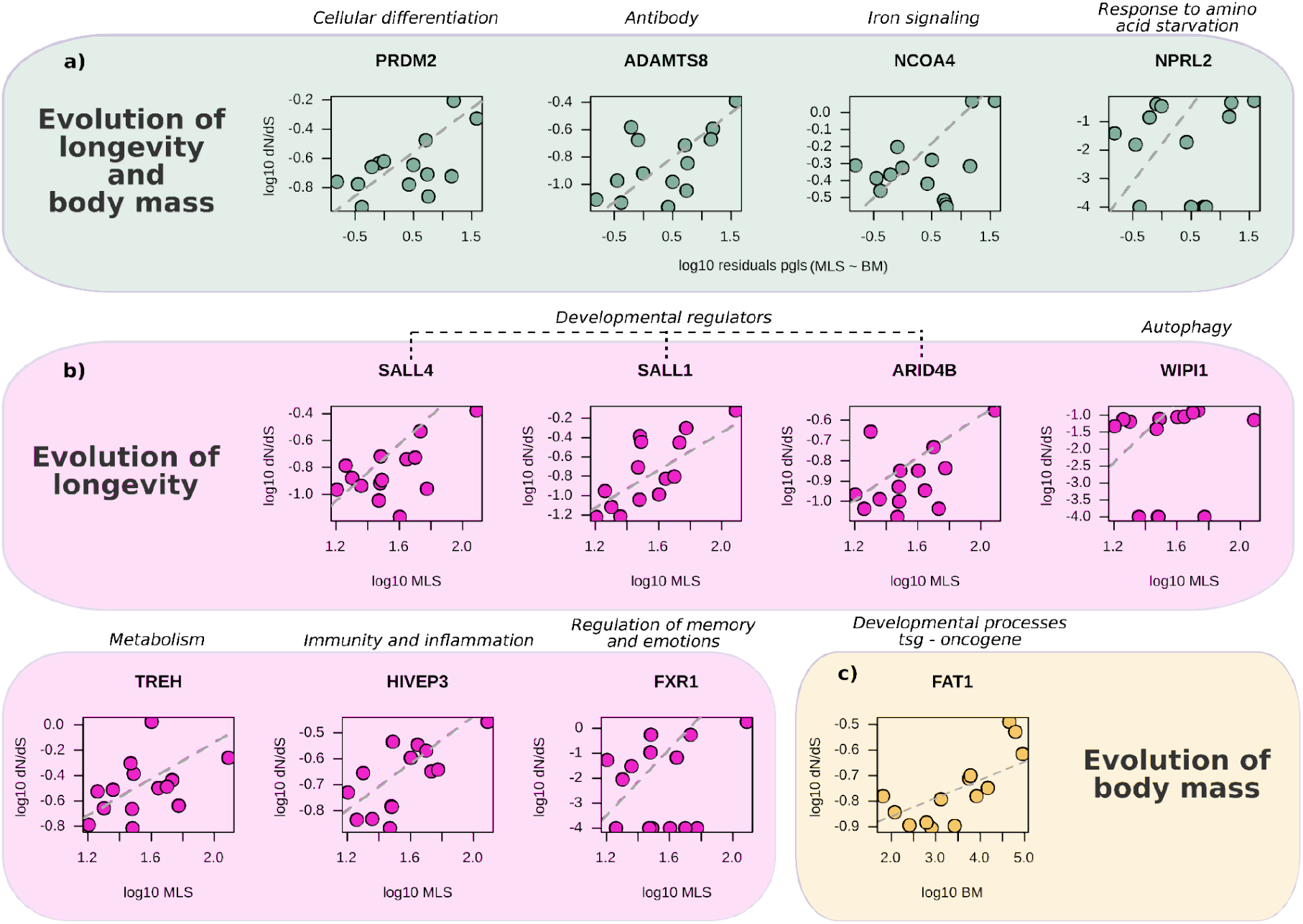
Examples of genes whose evolution correlated with the evolution of maximum lifespan (MLS) and/or body mass (BM). Three independent phylogenetic generalized least-squares models (PGLS) were performed in order to evaluate the independent contribution of each gene in the evolution of MLS and/or BM across primates. The dN/dS values were obtained from the branch model in PAML. a) PGLS model between dn/ds ~ residuals from the PGLS (MLS ~ BM). b) PGLS model between dn/ds ~ MLS and c) PGLS model between dn/ds ~ BM. In all the regression models the *p*-value was < 0.05 and the contribution of each gene (R^2^) > 22% (Table 1).

In the second PGLS model, we only focused exclusively on lifespan. We identified 58 genes that positively coevolved with longevity across primates (fig. 2b). Most of those genes encode for transcriptional regulators (Table S1.7) and are predominantly tumor suppressor or senescence genes. The genes *SALL1*, *SALL4* and *ARID4B* contribute from 37% to 53% of the variation in longevity seen across species (table 1b) and are important developmental regulators (Hirsch et al. 2015; Buttgereit et al. 2016; Wu et al. 2019; Bon-Baret et al. 2021; GÜven and Terzİ ÇİzmecİoĞlu 2021). Among the other genes, *WIPI1* is important for autophagy (Grimmel et al. 2015), *HIVEP3* for immunity and inflammation processes (Hicar et al. 2001; Krovi et al. 2020), and *FXR1* for the regulation of memory and emotions (Kalueff 2007; Sopova et al. 2019; Brown and Török 2021), important traits that have shaped primate evolution.

Finally, in the third PGLS model, we focused exclusively on body mass. This analysis revealed that *FAT1*, a highly conserved tumor suppressor gene involved in Wnt signaling pathway (Nusse and Varmus 1992), and associated with multiple cancers (Morris et al. 2013), contributes to ~22% of the variation in BM found across primates (table 1c).

In summary, we found that, across primates, the dN/dS ratio correlates with either longevity or body mass (or both) in a total of 67 genes. These genes are involved in multiple biological functions directly associated with ageing regulation, but are also related to particular features important for primate diversification such as diet, metabolism and neurological development.

### Longevity genes under positive selection in the ancestor of great apes are involved in neurological pathways, immunity and telomere length

The integration of an evolutionary perspective in the study of genes related to human health could reveal the discovery of novel molecular variants that underlie important biological questions, such as how to live longer and healthier lives, or how to be more resistant to diseases. Here, we studied the signal of positive selection in the evolution of 986 longevity genes in the ancestors of great apes, which is the lineage including the species with largest body sizes and longest lifespan across primates. We found six longevity genes with positive selection in the ancestors of great apes (*PAK1, ULK3, KIF1B, STAT1, ABCC3, TERF2;* fig. 3a-g; Table S1.8-1.9). These genes are involved in important biological functions that characterized the evolution of long-lived species, such as immunity, neurodevelopment, and telomere maintenance (Zhang et al. 2013; Gorbunova et al. 2014; Orkin et al. 2021). For example, *PAK1* is an oncogene involved in cytoskeletal reorganization and cortical development (Pan et al. 2015). The codons under positive selection are located in a protein kinase domain, indicating that these changes could have modified the structure or function of the gene in the ancestor of the great apes. Mutations in *PAK1* have been associated with macrocephaly in humans (Harms et al. 2018), and memory deficits in mice (Meng et al. 2005). In fact, this gene plays an important role in hippocampal synaptic plasticity (Asrar et al. 2009). Like *PAK1*, *ULK3* is also an important component of several neurological pathways, whose activity is significantly decreased in patients with Parkinson’s disease (Miki et al. 2018). KIF1B is an axonal precursor which encodes for an important motor protein that transports mitochondria. In humans, *KIF1B* mutations have been associated with autoimmune and neurological diseases, including multiple sclerosis (Aulchenko et al. 2008) and Charcot-Marie-Tooth (Zhao et al. 2001). *KIF1B*-knockout mice do not survive after birth due to atrophies in the nervous system, resembling what is observed in Charcot-Marie-Tooth patients (Zhao et al. 2001). *STAT1* regulates innate immunity and protection against viral infection (Karst et al. 2003; Hossain et al. 2019). *ABCC3* has been implicated in drug resistance, including chemotherapy (Balaji et al. 2016). Finally, *TERF2* is very important for ageing-related processes, since it regulates telomere length (Ilyenko et al. 2011; Tian et al. 2019). For this gene, the great apes show a specific substitution in the HTH myb-type Domain, a highly conserved amino-terminal region (Peters et al. 1987), suggesting that the substitution could potentially have affected the myb-type/DNA interaction important for TERF2 protein function.

**Figure 3.**
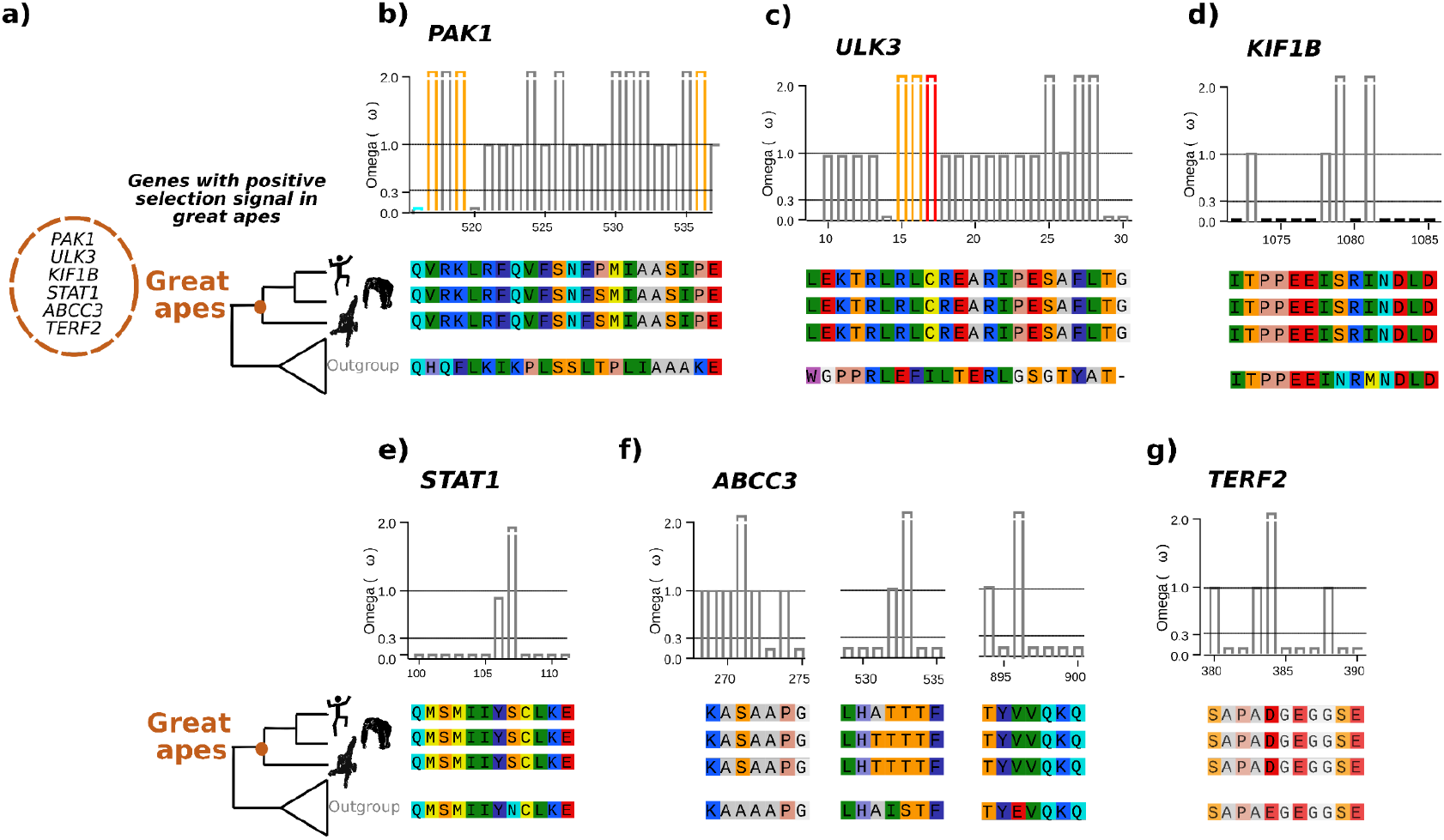
Genes under positive selection in the ancestor of great apes (a-g): *PAK1, ULK3, KIF1B, STAT1, ABCC3, TERF2*. The three first codon alignments on each gene represent fragments with an omega ω value >1. The colour gradients are used to display the degree and typology of amino acid change: the grey colour corresponds to the most conservative change, while red indicates the most radical. The alignments underneath display the consensus sequences across the outgroups (non-ape primates and other mammals). The amino acid colours reflect the different physicochemical properties.

### Longevity genes with human-specific expression patterns

We reasoned that different strategies may have contributed to the evolution of longevity genes in great apes. On the one hand, positive selection may lead to important changes in the amino acid composition, and thus in the structure and function of the encoded protein. On the other hand, the coding sequence, and hence the structure of the protein, may remain unaltered across species but the expression level of the gene may change. Changes in gene expression would be reflected in the amount of translated protein, which could have a significant impact on cell metabolism. Our data so far demonstrated that a number of longevity genes are under positive selection in great apes. We surmised that changes in the expression of longevity genes may provide an additional contribution to the differences in lifespan and size between the great apes and the other primates. To test this hypothesis, we leveraged a publicly available RNA-seq dataset generated from the liver of the six primate species analyzed in the present study (humans, chimpanzee, rhesus macaque, marmoset, mouse lemur and bushbaby; Trizzino et al. 2017). The liver is particularly relevant for this study because most genes associated with longevity and body mass are highly expressed in the liver, and also because liver gene expression displays high variation across species, likely reflecting adaptation to different diets and environments, in spite of a conserved core function (Trizzino et al. 2017). Consistent with this premise, the liver has been employed in studies that investigated biomarkers of aging (Lee et al. 2012; Bochkis et al. 2014; White et al. 2015).

As a first step, we examined the expression patterns of all the 1,553 six-way ortholog longevity genes, and found that genes involved in both ageing & senescence are significantly more expressed in humans relative to all the other primates grouped together (Wilcoxon rank-sum test, *p* < 0.001; fig. 4a). Similarly, genes that are involved in both ageing and tumor suppression are more expressed in humans than in other primates grouped together, although the *p*-value was only marginally significant, possibly as a consequence of the small sample size (N=3; Wilcoxon rank-sum test, *p* = 0.057; fig. 4b).

**Figure 4.**
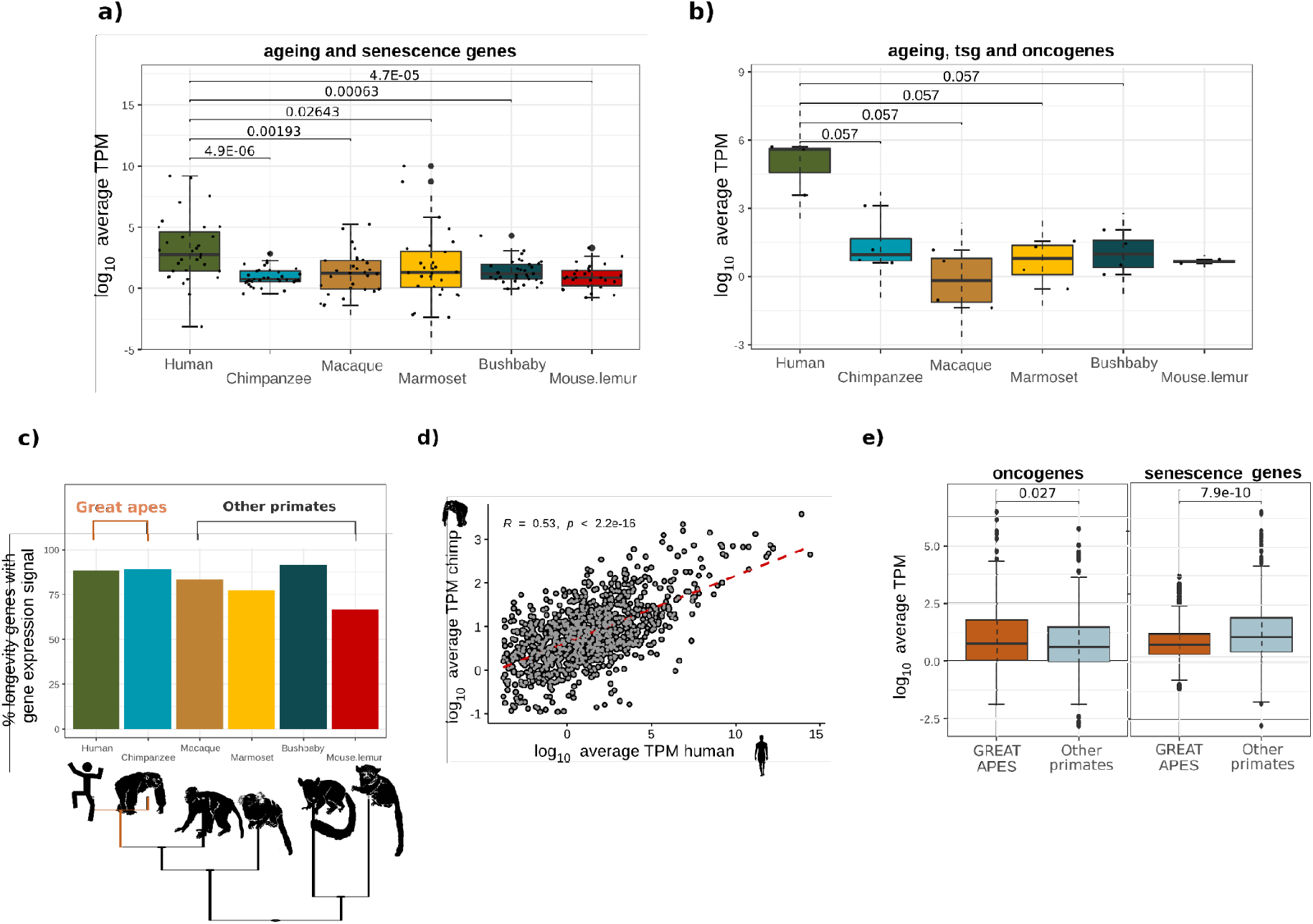
Evolution of longevity gene expression in the primate liver. Expression levels across species for the 1,553 longevity genes are divided into categories: ageing, senescence related genes, tumor suppressor genes (TSGs) and oncogenes; (a) ageing related genes and senescence genes (b) ageing related genes, tumor suppressors and oncogenes. In both groups, humans display higher expression compared to other primates (Wilcoxon rank-sum test, *p* < or near 0.05). c) The percentage of longevity genes expressed in the liver is similar across the six primate species. d) The expression of the 1,553 longevity genes in the liver is positively correlated between humans and chimpanzee (Spearman’s correlation coefficient, R=0.53, *p* < 0.001). e) Oncogenes are significantly more expressed in great apes compared to other primates. Senescence genes are significantly less expressed in great apes relative to the other primates (Wilcoxon rank-sum test, *p* < 0.05).

### Longevity genes with ape-specific expression patterns

Next, we compared the expression of the longevity genes between apes and other primates. Overall, apes have a higher number of longevity genes expressed in the liver relative to other primates (fig. 4c; Spearman’s correlation coefficient R=0.53, *p*-value = 2.2e-16), and the expression level of the longevity genes is strongly correlated between humans and chimpanzees (fig. 4d). We decomposed the longevity genes across the different categories (fig. 4e), and observed that the oncogenes are significantly more expressed in apes relative to other primates grouped together (Wilcoxon rank-sum test, *p* = 0.027), while the senescence genes are significantly less expressed in apes (Wilcoxon rank-sum test, *p* = 7.9 x 10^−10^). This pattern could reflect the findings from recent studies that reported that senescence genes are beneficial in the younger ages, during which they increase reproductive fitness, while they might have a negative impact later in life (Campisi 2003; Blagosklonny 2010; Di Micco et al. 2021). Accordingly, gene expression meta-analysis across human tissues showed that cancer genes promote longevity, whereas senescence genes showed anti-longevity patterns (Chatsirisupachai et al. 2019).

### Ape-specific enhancers drive the differential expression of longevity genes in apes

We performed a differential gene expression analysis, comparing the expression of the longevity genes in apes (human, chimpanzee) vs the other 4 primate species grouped together (rhesus macaque, marmoset, bushbaby, mouse lemur). Overall, 122/1,553 (~7.9%) longevity genes were found to be differentially expressed between apes and the other primates grouped together (DESeq2 FDR < 5%; Supplemental Table S3). Of those, 61 genes were upregulated and 61 downregulated. We investigated if cis-regulatory evolution could underlie the changes in expression of the 122 genes identified as differentially expressed in apes compared to other primates. To this purpose, we leveraged H3K27ac ChIP-seq data generated in the same study from the same liver samples of the same individuals (Trizzino et al. 2017). Namely, this specific histone modifications marks active cis-regulatory elements (enhancers and promoters). Importantly, in the original study, Trizzino and collaborators (2017) identified a set of ape-specific enhancers. We thus scanned the surrounding regions of the 122 differentially expressed genes found in the apes vs other primates comparison (fig. 5a). We focused on the 50 kilo-bases (kbs) up- down-stream of the transcription start site (TSS) of the 122 genes, and observed that 27 of the 122 differentially expressed genes (22.1%) have an ape-specific enhancer in the 50 kbs surrounding the TSS (38 total enhancers, median distance from TSS = 19.8 kb; fig. 5a; Table S3). To test if this number is higher than expected by chance, we performed a permutation test (1000 permutations), by randomly extracting (for 1000 times) 122 genes from a list including all the annotated human genes (Ensembl), and found that, on average, only 4/122 random genes (3.3%) were located within 50 kbs from an ape-specific enhancer. Together, these data indicate that the longevity genes identified as differentially expressed in apes vs other primates are ~six times more likely than expected to be found near an ape-specific enhancer (22.1% vs 3.3%; Permutation Test *p* < 2.2 x 10^−16^; fig. 5a).

**Figure 5.**
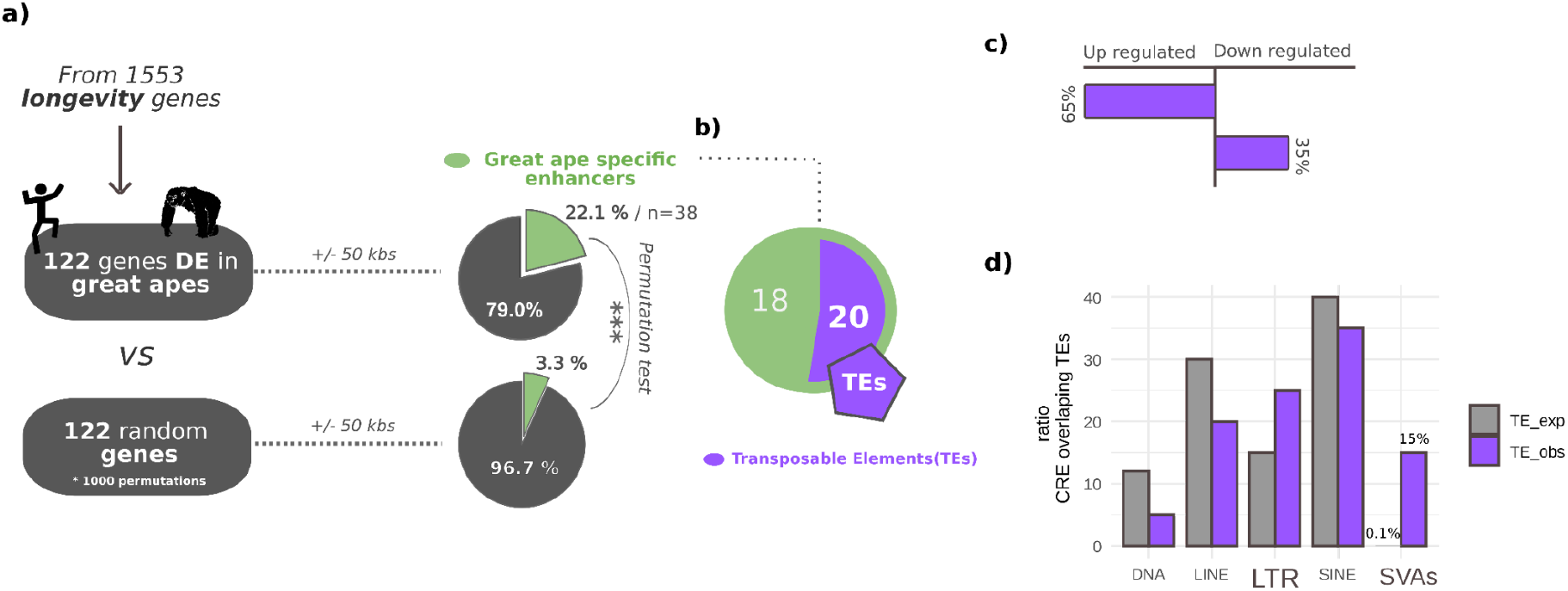
Ape-specific enhancers are enriched near differentially expressed longevity genes. a) 27/122 differentially expressed (DE) longevity genes (22.1%) are located within 50 kbs of an ape-specific enhancer (a total of 38 ape-specific enhancers for 27 DE genes). In comparison, only 4/122 (3.3%) randomly selected genes (1000 permutations) are located within 50 kbs of an ape-specific enhancer (Permutation Test *p* < 2.2 x 10-16).). b) 20/38 ape-specific enhancers (52%) located within 50 kbs of the DE longevity genes overlap annotated Transposable Elements (TEs). c) Of the DE longevity genes found near TE-derived ape-specific enhancers, 65% were identified as upregulated, 35% as downregulated (Fisher’s Exact Test *p* = 0.0425). e) LTR and SVA transposons are overrepresented in the sequence of the ape-specific enhancers located within 50 kbs of a DE longevity gene (Fisher’s Exact Test *p* < 2.2 x10^−16^ for SVAs; *p* < 0.0001 for LTRs).

### Ape-specific enhancers associated with differentially expressed longevity genes are derived from recent Transposable Element insertions

Overall, a total of 38 ape-specific enhancers were found in the 50 kbs surrounding 27 of the 122 differentially expressed longevity genes (Table S1.10). Since multiple studies have demonstrated that Transposable Elements (TEs) are an important source of gene regulatory novelty in the primate gene regulation (Jacques et al. 2013; Chuong et al. 2016; Trizzino et al. 2017; Trizzino et al. 2018), we tested for potential association between the 38 ape-specific enhancers and TE insertions. We found that 20/38 (~52.6%) ape-specific enhancers are derived from TE insertions (fig. 5b). This was not higher than expected by chance, considering that TEs represent ~50% of the human genome. Nonetheless, TE-derived ape-specific enhancers were significantly more associated with upregulated (65%) than to downregulated (35%) longevity genes (fig. 5c, one-tailed Fisher’s Exact Test *p* = 0.0425). Notably, *ADAMTS8,* a gene reported as under positive selection in other long-lived species (whales; Tejada-Martinez et al. 2021), and which in this study we identified as correlated with lifespan (fig. 2a), is upregulated in apes and is located near a TE-derived ape-specific enhancer.

Finally, we investigated if specific TE families were overrepresented in the TE-derived ape-specific enhancers associated with differentially expressed longevity genes (fig. 5d). Notably, 15% of those were derived from SINE-Vntr-Alu (SVA) insertions. SVAs are the youngest TE family, with most copies being either ape- or human-specific, and represent only 0.1% of all human (and 0.3% of chimpanzee’s) annotated TEs. Overall, our data suggest that SVAs are significantly overrepresented in our set of ape-specific enhancers associated with differentially expressed longevity genes (expected: 0.1% - observed 15%; Fisher’s Exact Test *p* < 2.2 x10^−16^). These findings are consistent with recent studies which demonstrated that SVA insertions contribute to human gene-regulatory networks (Trizzino et al. 2017; Trizzino et al. 2018; Pontis et al. 2019). Similarly, the LTR transposons were also overrepresented in our set of ape-specific enhancers (Fisher’s Exact Test *p* < 0.0001). These findings are also consistent with recent literature on the contribution of LTRs to human gene regulation (Wang et al. 2007; Cohen et al. 2009; Sundaram et al. 2014; Chuong et al. 2016; Janoušek et al. 2016; Fuentes 2018).

In summary our analysis revealed that lineage specific enhancers may have contributed to the evolution of the expression of longevity genes in great apes and that young transposon insertions may have had a significant role in this process.

## Discussion

Natural selection and gene regulatory evolution likely underlay the evolution of most phenotypic traits. Here, we specifically focused on the evolution of lifespan and body mass in great apes. This primate lineage includes species which evolved with a lifespan and body mass significantly different from the rest of primates. We carried out comparative genomic analyses, examining different sources of molecular variation, including both coding genes and non-coding genomic regions (i.e. cis-regulatory elements: enhancers, and promoters). Cis-regulatory evolution plays an essential role in phenotypic diversification (Trizzino et al. 2017; Berthelot et al. 2018; Farré et al. 2019; Feigin et al. 2019; Sundaram and Wysocka 2020; Marand et al. 2021), and can lead to evolutionary innovations in disease resistance across species (Gorbunova et al. 2007; MacRae et al. 2015; Tollis et al. 2020).

Our comparative genomic analysis revealed that 67 genes positively correlated with the evolution of maximum lifespan and body mass in primates. Out of those, 85% co-evolved exclusively with maximum lifespan, while the remaining 15% co-evolved with lifespan and body mass together, or with body mass alone. These genes are mostly involved in tumor suppression and senescence-related processes. Since the increase in longevity elicits a greater risk to develop cancer, the rise and fixation of new molecular variants (non-synonymous substitutions) could have positively contributed to improve the functionality of the tumor suppressor genes and thus be beneficial for the evolution of the most recent primate lineage. Similar outcomes have been recently reported by a study on mammals focused on copy number variation in tumor suppressor genes (Sulak et al. 2016; Tollis et al. 2020). Consistent with these lines of evidence, previous studies have found that the rate of protein evolution in anti-cancer and DNA damage response genes is accelerated in long-lived mammals (Li and de Magalhães 2013; Tollis et al. 2019; Tejada-Martinez et al. 2021).

A greater lifespan correlates with a higher likelihood of being exposed to pathogens and diseases. Consequently, the immune system of long-lived species is expected to be under particularly strong pressure to sustain the prolonged arm-race with the pathogens (Quesada et al. 2019; Singh et al. 2019). Consistent with this premise, we found a significant number of immune, inflammation, and DNA repair genes that coevolved with longevity in primates. Notably, several of those genes were also under positive selection in the ancestors of great apes. This points towards an evolutionary strategy in which disease resistance and lifespan evolved together in primates, and particularly in apes, as it has been reported for other long-lived species (Harris et al. 2017; Muntané et al. 2018; Vazquez et al. 2018).

In the great ape ancestor, we identified genes under positive selection involved in both the immune system and in the regulation of telomere length. Importantly, telomeres play a key role in ageing-related processes, and the regulation of telomere length is an important mechanism to control cancer in species with increases in body mass (Seluanov et al. 2008). Namely, telomeres shorten with age, and this leads to increased frequency of chromosomal damage, cell senescence and apoptosis (Tian et al. 2018). It has been proposed that in large-size and long-lived species the rate of telomere shortening is slower than in smaller and shorter-lived species (Whittemore et al. 2019). Based on our data, we speculate that genes involved in telomere function and length may have evolved to help great apes fight diseases, ultimately contributing to their longer lifespan.

Among the genes with positive selections in apes, we identified several associated with diet, metabolism, as well as with the regulation of memory and emotions, all of which are important great ape features. This finding is consistent with several studies that have proposed a link between increased life span and increased cognitive functions (Ghirlanda et al. 2014; Barton and Venditti 2017; Orkin et al. 2021). Characterizing their molecular evolution could help us to better understand the rise of genetic neurological disorders in humans, such as macrocephaly, cognitive disorders and Parkinson’s disease. Different molecular strategies may have contributed to the evolution of longevity genes across primates. Positive selection can lead to changes in the structure and function of a protein through amino acid substitutions, without affecting gene expression, and thus the amount of protein produced by a specific cell-type. In parallel, the expression level of genes important for longevity and body mass could be affected by evolutionary changes in the associated gene regulatory network. For example, a novel, species-specific, enhancer could influence the expression of the associated gene, ultimately leading to higher levels of protein production without any modification to the structure of the protein itself. Here, using RNA-seq we have investigated how the expression of the longevity genes varies across primates, using the liver as a proof of principle. We demonstrated that oncogenes are significantly more expressed, and senescence genes less expressed, in great apes relative to other primates, suggesting a tradeoff between living longer (which leads to increased likelihood to produce offspring) and disease susceptibility later in life (Campisi 2003; Blagosklonny 2010; Rodríguez et al. 2017; Di Micco et al. 2021).

Overall, we report that ~8% of the longevity genes tested in this work were differentially expressed in the liver of great apes relative to higher primates, and we demonstrate that lineage specific (i.e. ape-specific) enhancers have significantly contributed to this process. In fact, longevity genes differentially expressed between great apes and other primates are ~six times more likely than expected to be located near an ape-specific enhancer. Consistent with recent studies that suggested an important contribution of young transposable elements (TEs) in the evolution of primate gene regulation (Jacques et al. 2013; Trizzino et al. 2017; Pontis et al. 2019), we demonstrate that SVA transposons are significantly overrepresented in the DNA sequence of ape-specific enhancers located near differentially expressed longevity genes. The SVAs are the youngest family of TEs, and include ape- and/or human specific copies.

In summary, our work has shed light on the evolution of thousands of longevity genes in great apes, highlighting a significant contribution to this process for both positive selection on coding genes, as well as evolutionary changes in the non-coding regulatory regions.

## Materials and Methods

### Relationship between body mass and longevity across primates

To determine if the body mass contributes to the variation in longevity independently in primates relative to other mammals, and to assess ape-specific patterns, we examined the evolution of the life history traits maximum lifespan (MLS) and adult body mass (BM) across 932 mammals. We fit a linear regression through the phylogenetic generalized least squares (PGLS; (Orme et al. 2012), while controlling for potential phylogenetic signals. Two independent regression models were implemented: the first one across primates, and in the second model, the regression was performed between great apes vs other primates (log10 MLS ~ log10 BM * great_apes). The phylogenetic tree used was derived from Uyeda et al. (2017) (Uyeda et al. 2017) and the MLS and BM data were gathered from AnAge database (Tacutu et al. 2018).

### Genomic sampling, longevity associated genes and build 2 of CellAge Database of Cell Senescence Genes

To study the evolution of longevity in primates we carried out a phylogenetic design that included genomes from 19 mammalian species. Our taxonomic sampling included 3 species from the family Hominidae (*Homo sapiens* - human, *Pan troglodytes* - chimpanzee and *Pongo abelii* - sumatran orangutan), one representant from Hylobatidae (*Nomascus leucogenys* - white-cheeked gibbon), three Cercopithecidae (*Macaca mulatta* - Rhesus macaque, *Rhinopithecus roxellana* - Golden snub-nosed monkey and *Chlorocebus sabaeus* - vervet), four Platyrrhini (*Callithrix jacchus* - white tufted ear marmoset, *Aotus nancymaae* - night monkey, *Saimiri boliviensis* - bolivian squirrel monkey, and *Cebus imitator* - white-faced capuchin), two Strepsirrhine (*Otolemur garnettii* - bushbaby and *Microcebus murinus* - Gray mouse lemur) and finally 5 other mammalian species outside of primates (*Tupaia chinensis -* Chinese tree shrew, *Mus musculus* - common mice, *Canis lupus familiaris* - dog, *Bos taurus* - cow and *Sus scrofa* - pig).

The coding sequences of each species were downloaded from Ensembl v.96 and NCBI databases (Table S1). To remove low quality records, sequences were clustered using CD-HIT-est v.4.6(Fu et al. 2012) with a sequence identity threshold of 90% and an alignment coverage control of 80%. The longest open reading frame was kept using TransDecoder LongOrfs and TransDecoder-predicted in TransDecoder v3.0.1 (https://github.com/TransDecoder/TransDecoder/).

The longevity associated coding genes, 2,268 in total, were gathered in four different categories: ageing genes from GenAge Database (build 20 - 307 protein coding genes, https://genomics.senescence.info/genes/index.html); TSGs from the Tumor Suppressor gene database (TSGene 2.0 - 1018 protein coding genes, https://bioinfo.uth.edu/TSGene/; (Zhao et al. 2013); oncogenes from the Oncogene database (698 protein coding genes, http://ongene.bioinfo-minzhao.org/; (Liu et al. 2017); and finally, we compiled build 2 of the CellAge Database of Cell Senescence Genes with an additional 655 genes, complementary to the 279 genes previously reported; https://genomics.senescence.info/cells/, (Avelar et al. 2020).

Build 2 of CellAge was compiled in the same way as build 1. A robust scientific literature search was performed for relevant papers before curation and annotation; genes were appended to the database if they met the following criteria:

· Only gene manipulation experiments were used to identify the role of the genes in CS to ensure objectivity in the selection process.
· The genetic manipulation caused cells to induce or inhibit the CS process in wet lab experiments. CS was detected by growth arrest, increased SA-β-galactosidase activity, SA-heterochromatin foci, a decrease in BrdU incorporation, changes in morphology, and/or specific gene expression signatures such as p21 and p16.
· The experiments were performed in primary, immortalized, or cancer human cell lines.

The resulting list comprised 934 genes that in human cell lines can induce or inhibit the CS process (Table S2). The full CellAge database, including all annotations regarding cell types and cell lines is available at https://genomics.senescence.info/cells/.

### Homology inference

We inferred homology relationships among the 2,268 longevity genes and the 19 mammalian species included in our study using the program OMA standalone *v*.2.3.1 (Roth et al. 2008; Altenhoff et al. 2019). We inferred the OMA Groups (OG), containing the sets of orthologous genes, for which we performed natural selection analyses. The amino acid sequences were aligned using the L-INS-i algorithm from MAFFT v.7 (Katoh and Standley 2013) and the codon alignments using the function pxaa2cdn in phyx (Brown et al. 2017). Finally, to reduce the chance of false positives given for low quality alignment regions we used the codon.clean.msa algorithm of the rphast package (Hubisz et al. 2011) with the associated longevity gene from the OG as reference sequence.

### Evolution of longevity genes and life history traits

To evaluate the possible co-evolution between the longevity genes and the life history traits MLS and BM in primates, the rate of evolution (ω=dN/dS) was calculated. The ω ratio per gene was estimated for each tip in the 14 primates species using the branch model (Yang 2007) as is implemented in ETE-toolkit with the *ete-evol* function (Huerta-Cepas et al. 2016). To calculate the ω ratios the treeshrew, dog, cow and pig were used as outgroups. Three independent PGLS models were performed per OG: 1) between the ω and the MLS; 2) between the ω and the BM; and 3) between the ω and the PGLS residuals between the MLS ~ BM. Previous to the analysis, all variables were transformed to log10. The genes with a dN/dS>2 were removed from the analysis.

### Positive selection analysis

To evaluate the role of natural selection in the evolution of longevity genes in the ancestor of great apes, we used codon-based models through a maximum likelihood approach using the program PAML v4.9 (Yang 2007), as is implemented in ETE-toolkit with the *ete-evol* function (Huerta-Cepas et al. 2016). We calculated the branch-site model in order to estimate changes in the ω value of individual sites. We compared the null model, where the ω value in the foreground branch was set to 1, with the model in which the ω value was estimated from the data (Zhang 2005). The comparisons between models were made using likelihood ratio tests (LRT) and the p-values from the LRT were corrected with FDR (Benjamini and Hochberg 1995). All the genes with positive selection signal but with a LRT > 1 were manually checked for false positives.

### RNA-seq analysis

To evaluate the contribution of the gene regulation to the evolution of longevity genes in great apes, we took advantage of recently published RNAseq and Chip-seq data for histone H3 lysine 27 acetylation (H3K27ac) from liver of six species representatives of the main groups of primates (human, chimpanzee, rhesus macaque, marmoset, bushbaby and gray mouse lemur; (Trizzino et al. 2017). The list of the genes differentially expressed between Apes and other primates was downloaded from the Supplementary Materials of the same paper (Trizzino et al. 2017). To generate box-plots comparing Gene Expression (GE) across species, we leveraged the transcripts per million (TPM), also available from the Supplementary Materials of Trizzino et al (2017). The TPMs were quantile normalized using R version 3.6.3 (R Core Team 2020).

### Enhancer evolution analysis

From the publicly available Chip-seq data (Trizzino et al. 2017), we downloaded the list and coordinates all the enhancers previously identified as “ape-specific” by Trizzino and collaborators. We then selected all the ape-specific enhancers overlapping a region encompassing +/− 50 kbs from the transcription start site (TSS) of each differentially expressed longevity gene using BEDTools v2.29.2 (Quinlan and Hall 2010). Based on this association, we then calculated how many longevity genes were both differentially expressed in the ape vs other primates comparison AND also associated with an ape-specific enhancer. To evaluate if this number was higher than expected by chance, we selected 122 random genes in the human genome and examined how many of them had an ape-specific enhancer within 50 kbs of the TSS and assessed statistical significance by means of Fisher’s exact test. Finally, using BEDTools, we looked for overlap between transposable elements and ape-specific enhancers associated with genes differentially expressed in ape vs primate comparison. The list and coordinates of the transposable elements were downloaded from the UCSC Genome Browser (RepeatMasker; hg38 assembly).

### Enrichment analysis

To gain insight into the particular functions of the longevity genes of interest, the enrichment analysis was performed using WebGestaltR v0.4.4 (Liao et al. 2019). We tested for significant pathway associations using the hypergeometric test for Over-Representation Analysis (ORA) (Khatri et al. 2012). We selected the categories of gene ontology, biological processes, molecular pathways, diseases OMIM, and human phenotype and we considered overrepresented categories to be those with a significance level above that of an FDR of 0.01 after correction with the Benjamini-Hochberg multiple test. An independent enrichment analysis was made using as a background the protein-coding genes and relative to the 2,268 longevity genes.

### Libraries in R associated with the data treatment, statistical analysis and graphics

Bioconductor, msa, ape, geiger, nlme, pythools, caper, ggpubr, gridExtra, tidyverse, grid, corrplot, ggplot2, ggpubr, Hmisc, gmodels, car, DescTools, qqplotr, dplyr, RColorBrewer, calecopal.

## Acknowledgements

This study was funded by NIH R35 GM138344-01, awarded to M.T. AnAge, GenAge and CellAge are supported by funding from the Biotechnology and Biological Sciences Research Council (BB/R014949/1) to J.P.M. The authors are grateful to Alejandra Tejada-Martinez for the graphical support.

## Contributions

D.T.M. and M.T. conceived the study and performed the genomic analyses. R.A., J.P.M., B.Z. and G.N. compiled the Build 2 of the CellAge database. I.L. contributed to the Cellage website development. D.T.M., M.T. and R.A. wrote the manuscript. M.T. and D.T.M. directed the research.

