## Supplementary_materials for "Positive selection and enhancer evolution shaped lifespan and body mass in great apes": Supplementary figure1.docx

**Supplementary figures**

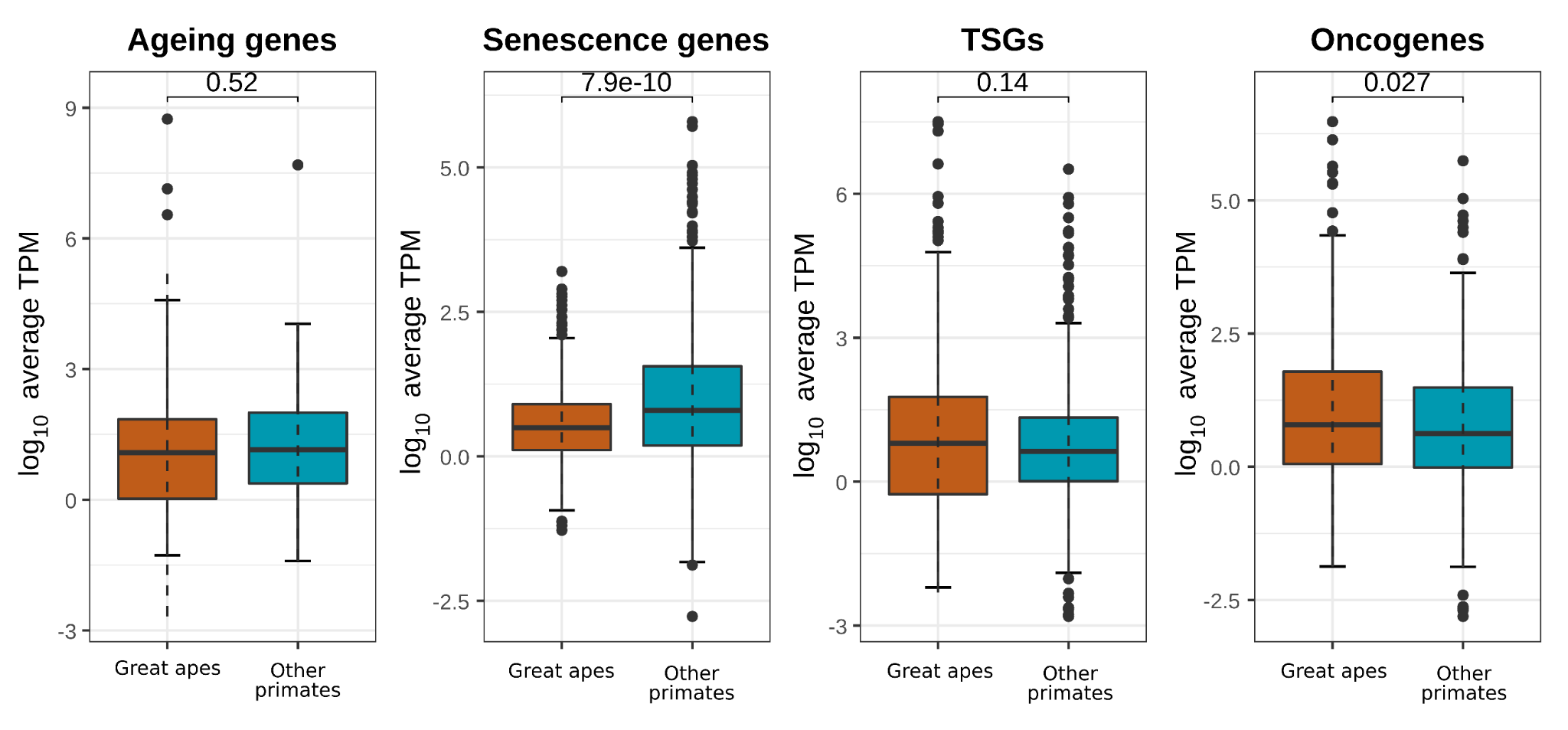

**Supplementary figure 1.** Expression levels between great apes vs other primates for the 1,553 longevity genes. Senescence genes are significantly less expressed in great apes relative to the other primates while Oncogenes are significantly more expressed (Wilcoxon rank-sum test, p-value < 0.05).
